# Gene expression signatures predict circadian rhythms in oncogenic pathways

**DOI:** 10.1101/2022.11.02.514919

**Authors:** Eleonora Winkler, Bharath Ananthasubramaniam, Hanspeter Herzel

## Abstract

Day-night environmental cycles together with our own adaptive rhythms in behavior and physiology lead to rhythmicity of various processes on the cellular level, including cell signaling. Despite many implications of such daily changes in signaling, the quantification of such rhythms and estimates of peak phases of pathway activities in various tissues are missing. Governed mainly by posttranslational modifications, a pathway activity might not be well quantified via the expression level of pathway components. Instead, a gene expression signatures approach can be used to score activity of various pathways. Here, we apply such gene expression signatures on circadian time series transcriptomics data to infer rhythmicity in cellular signaling. We show that, across multiple datasets, the gene expression signatures predict the presence of rhythmicity in EGFR, PI3K and p53 pathways in mouse liver. With the focus on EGFR pathway, we pinpoint the most influential signature genes for the overall rhythmicity in the activity scores for this pathway. These findings suggest that time of the day is an important factor to consider in studies on signaling. Simultaneously, this study provides a new paradigm to use circadian transcriptomics to get at temporal dynamics of pathway activation.

## I. Introduction

Mammalian biology is orchestrated along the 24 h day by means of an internal time-keeping system, called the circadian clock. A gene-regulatory network that forms the circadian clock produces cell-autonomous oscillations in the abundance of its components with a period close to 24 h (Kramer and Merrow, 2013; Takahashi, 2016). Rhythmic inputs to an organ (such as feeding) and the coupled cell-intrinsic clocks in the organ tissues can both drive daily oscillations in tissue transcriptome and proteome (Balsalobre et al., 2000; Guan et al., 2020; Koronowski et al., 2019). In most studied tissues, 10-40 % of transcripts and 6-20% of cellular proteins oscillate, affecting most cellular functions (Robles et al., 2014; Wang et al., 2017; Zhang et al., 2014). Notably, there is accumulating evidence that many signaling pathways are rhythmically activated in anticipation of or in response to daily environmental changes (Aviram et al., 2021; Goldsmith et al., 2018; Horiguchi et al., 2013; Vollmers et al., 2009).

Systems biology tools, such as set enrichment analysis, applied to the transcriptomic time series data have shed light on the functional relevance of the rhythmicity in the transcriptome including signaling pathway activities. One such common approach looks at enrichment in the signaling pathway components among rhythmic transcripts with similar peak phases (Zhang et al., 2016). In some studies, the structure of the pathway, i.e. whether certain pathway components have opposing roles, is also taken into account (Acevedo et al., 2021; Zhang et al., 2014). However, evaluating the overrepresentation of pathway components among the rhythmic genes can still be misleading, since the circadian control of just a few rate-limiting steps might be sufficient to induce rhythms in a pathway function (Fustin et al., 2012; Kim and Reed, 2021; Panda et al., 2002).

Another important assumption behind the application of gene set enrichment methods on transcriptomics data is the correlation of pathway gene expression with pathway activity. However, it has been shown that expression of pathway components is not a reliable proxy for pathway activity due to the crucial importance of the posttranslational regulation (e.g. protein phosphorylation and ubiquitylation) (Buccitelli and Selbach, 2020; Nusinow et al., 2020; Robles et al., 2017; Wang et al., 2018). Thus, in the field of cell signaling research, alternative approaches are being explored, including the gene expression signature approach (Watters and Roberts, 2006). In the latter, consistent gene expression “footprints” of pathway activation/inhibition are derived from pathway perturbation experiments and then used to score the pathway activity in a transcriptomics dataset or to compare scores between samples (e.g., normal vs. tumor).

Gene expression signatures are usually used to infer pathway activation from static gene expression data. Here, we employ this approach to reconstruct dynamic pathway activation in the context of circadian rhythms. For that purpose, we use PROGENy software as it is applicable to mouse datasets and was designed in an attempt to capture universal (shared among different tissues and conditions) primary responses to perturbations of multiple cancer-related signaling pathways (Holland et al., 2020; Schubert et al., 2018). We quantify pathway activities at different time points in time series datasets and then search for daily rhythmicity in the resulting pathway activity scores.

We show that mouse liver transcriptomics data exhibit daily oscillations in the pathway activity scores for many pathways. Notably, many genes that are known to be rhythmic *in vivo*, or even *in vitro* (core clock genes), are present in the signature gene sets. With the focus on the epidermal growth factor receptor (EGFR) pathway, a crucial pathway involved in cell proliferation and survival, we show which genes drive the rhythmicity of the PROGENy scores and compare these for different time series datasets. Finally, we summarize the possible sources of daily rhythms in the signaling network downstream of EGFR and related receptors.

## II. Results

### Pipeline design

We apply the gene expression signature approach to the transcriptomics time series data to search for the “rhythmic footprints” of signaling pathway activations in mouse liver. We use the PROGENy software to score the activity of 14 cancer-related pathways in each sample, so that a time series collection of samples produces a time series of pathway activity scores (Fig. 1). We then analyze if there is statistically significant rhythmicity in the scores (24 h period, RAIN software with q-value cutoff 0.05).

**Fig.1.**
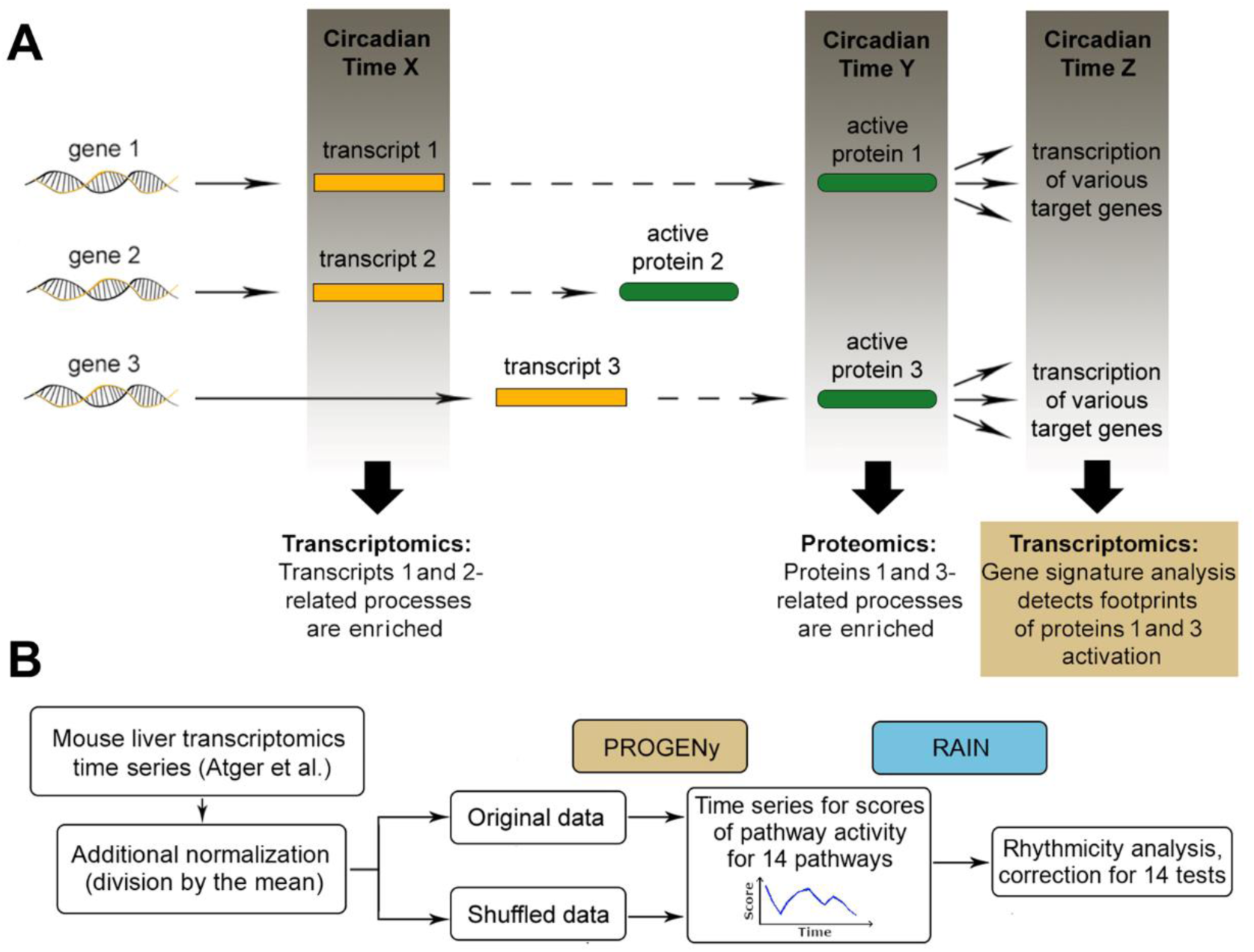
Gene expression signature approach applied to time series circadian data. (A) Different approaches to infer rhythmicity of a process/pathway via omics data analysis: set enrichment analysis for pathway components vs. gene expression signature approach that focuses on target genes. (B) The main pipeline of this study consists of PROGENy score calculation for time series data plus rhythmicity analysis of the score with RAIN.

The PROGENy score for a pathway and a given sample is a sum of weighted expression levels for all signature genes. Thus, highly expressed genes dominate the signal, especially when they have a high absolute weight coefficient. However, our goal was to discern the rhythmic signal in the data, even if it is present in the lowly expressed genes. In other words, a highly rhythmic gene that has a low mean level is still of interest since it can represent a “footprint” of rhythmic pathway activation. Thus, we added an additional normalization step to the transcriptomics data processing, where gene expression values were divided by the mean expression value for every gene (see Fig.S1A for results without this normalization step).

### Mouse liver gene expression suggests rhythmic activation of multiple pathways

We applied the approach described above to the circadian transcriptomics dataset from Atger et al. (Atger et al., 2015). Out of 14 pathways, 9 were considered significantly rhythmic according to our criteria (Fig.2A and S2). For comparison, we repeatedly shuffled the time points for each gene, destroying the temporal pattern in gene expression in the data. We observed that in shuffled datasets there were on average no rhythmic pathways. Treating RAIN q-values as a “rhythmicity measure” in the data, we can compare the distribution of the q-values after shuffling with the original q-values from the Atger dataset (Fig.2B). Alternatively, amplitudes of the absolute scores’ oscillations can be compared (Fig.S1B). Fig.S1C shows that the selection of an appropriate background model is essential.

**Fig.2.**
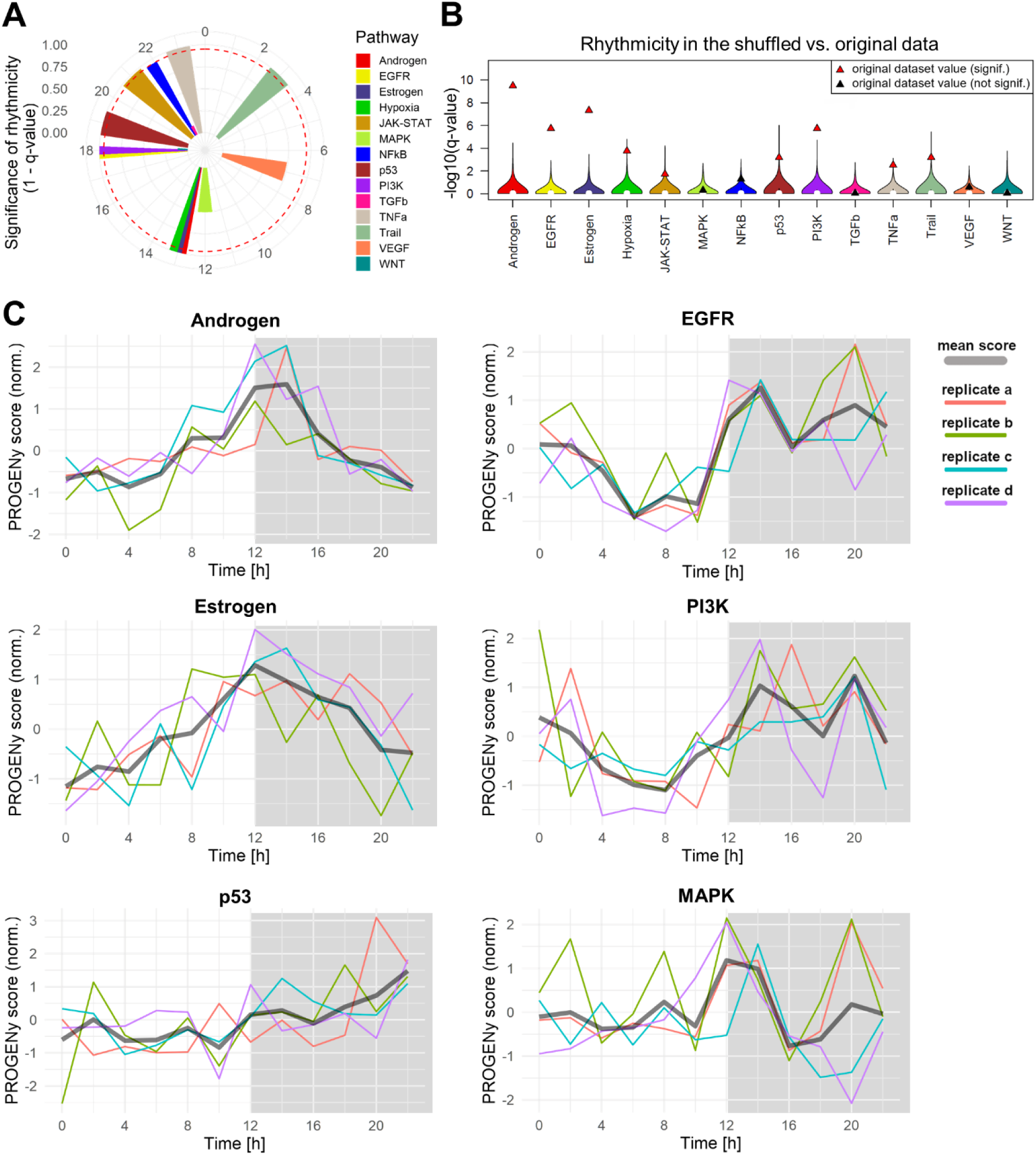
Most pathways exhibit daily rhythms in PROGENy scores (Atger et al. dataset). **(A)** Peak phases of PROGENy scores for all 14 pathways; height of the bars corresponds to significance of rhythmicity (1 – q-value) and the dashed red line designates the significance cutoff. **(B)** Distributions of the q-values acquired from shuffled data (times shuffled for each gene) in comparison with original q-values. **(C)** PROGENy scores time series for the four most significantly rhythmic pathways, as well as p53 and MAPK pathways; colors correspond to individual replicates, labeled as in the original dataset.

The lowest q-values were calculated for androgen and estrogen pathways, both peaking at around CT13, i.e., at the beginning of subjective night (Fig.2C). The same peak phase of these endocrine pathways might stem from the fact that estrogens in male mice are generated solely from androgens, and thus might follow the same pattern of rhythmicity. The next two pathways with most significant rhythms were EGFR and PI3K, with highest scores during the night (CT14-20). EGFR and PI3K pathways are activated downstream of various receptor tyrosine kinases and exhibit extensive crosstalk. Another pathway related to EGFR, namely the MAPK pathway, was not significantly rhythmic. Nevertheless, MAPK score also follows a pattern similar to EGFR and PI3K pathways.

To explore whether multiple signature genes are responsible for the rhythmicity in the scores, we employed a jackknife-like approach. Signature genes were removed from the dataset one at a time, and the resulting dataset was sent through the original data analysis pipeline. For all significantly rhythmic pathways, removal of any one gene did not abolish rhythmicity, highlighting that multiple signature genes are responsible for the observed rhythmic signals (Fig.S3).

### EGFR pathway: rhythmic contributions of individual genes to the scores tend to be in phase with each other

Motivated by other studies (see Discussion) and our own results on daily rhythmicity in the EGFR pathway activation, we focused on EGFR pathway for further analysis. EGFR is a receptor tyrosine kinase that is activated by various growth factors and contributes to the regulation of cell survival and proliferation. To better understand the PROGENy EGFR score rhythm, we looked at the individual contributions of different genes to the score, with a focus on genes with a 24 h period rhythmicity. Such rhythmic genes in the dataset were determined with RAIN software (FDR of 5%) and filtered for a peak-to-mean fold-change of at least 1.2. How strongly a signature gene contributes to the final score depends only on the relative amplitude and the PROGENy weight for this signature gene. In Fig.3A, relative amplitude multiplied by the weight coefficient is plotted against the peak phase of a gene. Strikingly, genes with negative PROGENy weight (i.e., driving the score down) all peak during the day, when EGFR score is lowest. In contrast, almost all genes with positive weight (i.e., driving the score up) peak during the night, when the EGFR score reaches its peak. Several genes stand out as most influential: Nr1d2 (a circadian clock component), Gabarapl1, Odc1, Lipg, and Phlda2.

**Fig.3.**
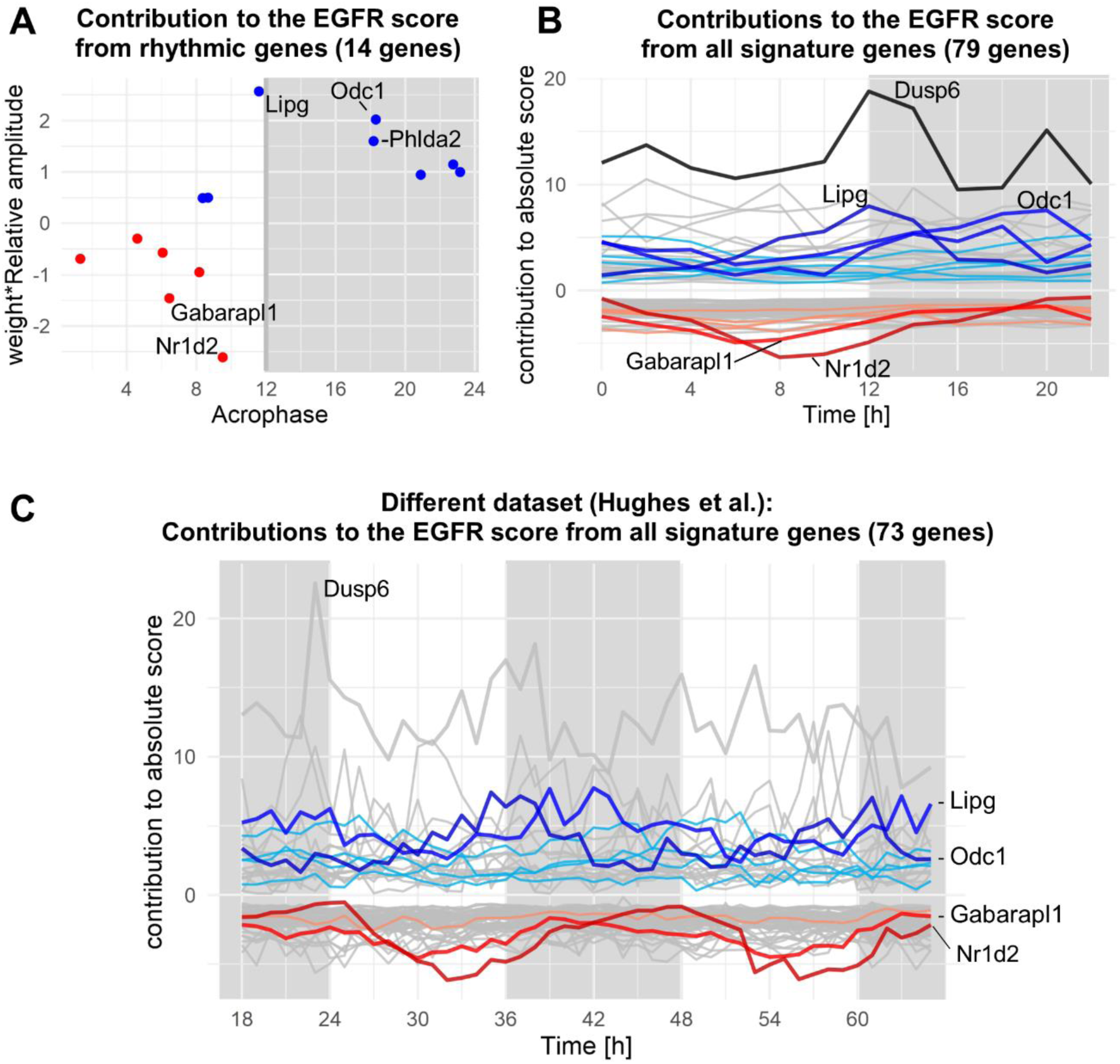
Contributions of rhythmic genes to the EGFR score tend to be in phase with one another. **(A)** Peak phases of rhythmic signature genes plotted against the strength of their contribution to the score (for Atger et al. dataset). Color of the dots corresponds to the sign of the weight coefficient. Genes with most influence on the score are the highest (positive weight) and lowest (negative weight) dots on the graph, their names are indicated. **(B)** Contributions to the score from all the signature genes present in the Atger dataset plotted as time series. Rhythmic genes are depicted with colors (blue for positive weight genes, red for negative), and the most influential genes are highlighted with stronger colors. Dusp6, not significantly rhythmic in the dataset, is highlighted due to its high PROGENy weight. In **(A)** and **(B)** replicates are averaged, grey areas on the graphs correspond to subjective night hours. **(C)** Contributions to the score from all the signature genes in a different dataset (Hughes et al.), plotted analogously to **(B)**.

To check if some genes that were not recognized as rhythmic still had considerable influence on the shape of the score time-series, we plotted contributions to the score of all signature genes at all time points (Fig.3B). Indeed, Dusp6, although not significantly rhythmic in mouse liver according to our analysis and previous reports (Tsuchiya et al., 2013), appeared to be influential due to a large PROGENy weight and a temporally non-homogeneous expression: a primary peak at CT12-14 and a secondary peak at CT20 with a highly variably expression between replicates. Since expression values are multiplied by the weight coefficient, this pattern is further magnified and is reflected in the overall EGFR score.

Overall, this analysis allowed us to “deconstruct” the EGFR score and pinpoint the genes most influential for the score rhythmicity and the overall shape. We then removed all the rhythmic genes from the dataset and observed how the rhythmicity q-values and score amplitudes change. Notably, without all rhythmic signature genes the score remains borderline rhythmic, probably due to the cumulative effect of the remaining low-amplitude contributions from other signature genes (Fig.S4). For comparison, Fig.3C shows a similar decomposition of the EGFR score for a dataset from Hughes et al., which is further discussed in the next section.

### EGFR rhythms are consistently observed across multiple mouse liver datasets

As a consistency check, we applied our pipeline to several other mouse liver datasets, with results summarized in Fig.4A. The Atger dataset includes data obtained from mice, subjected to restricted feeding, which produced results comparable to the ad libitum fed mice. Analogously, in two commonly studied microarray datasets from Hughes et al. (Fig.3C and S5) and Zhang et al., mouse liver samples produced consistent results for the EGFR pathway (Hughes et al., 2009; Zhang et al., 2014). Notably, across these datasets there is a high overlap of genes that are most influential to the score rhythmicity (Fig.4B).

**Fig.4.**
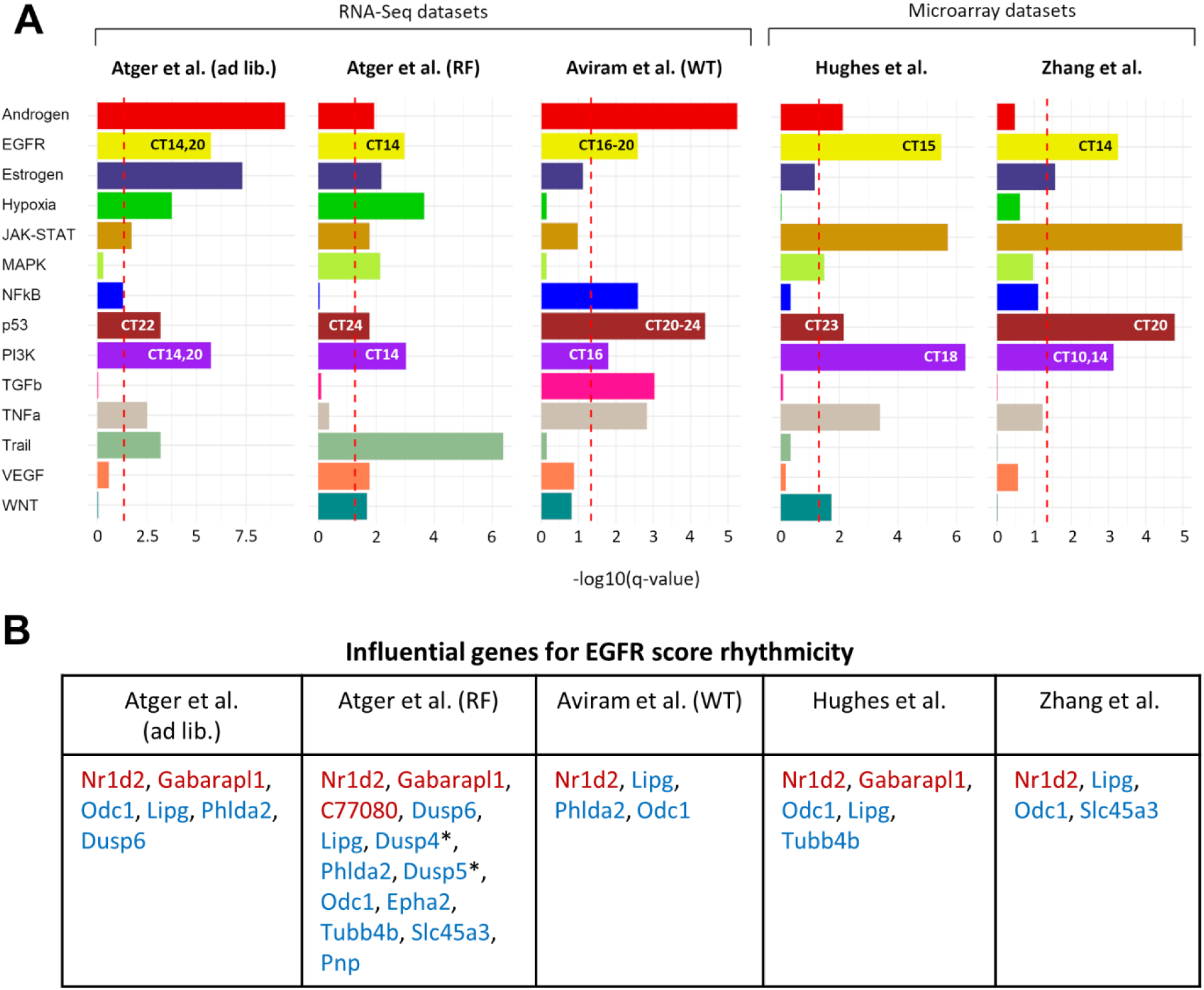
Multiple mouse liver datasets share similar score rhythms for EGFR, PI3K and p53. **(A)** Comparison of rhythmicity q-values across datasets; the dashed red line designates the significance cutoff. Peak phases for EGFR, PI3K and p53 pathways are indicated on the respective bars. **(B)** Negative (red) and positive-weighted (blue) genes that are influential for the EGFR score rhythmicity due to the large relative amplitude and absolute weight. Asterisks (*) mark the genes that have score contribution out of phase with the final score (between CT1 and CT10). Ad lib. – ad libitum feeding; RF – restricted feeding.

Apart from EGFR, we have noticed that PI3K and p53 also remain consistently rhythmic in all the depicted datasets. A deeper look into those pathways can be found in the supplement (Fig.S6). Overall, daily rhythms in the PROGENy EGFR, as well as PI3K and p53, scores can be observed in mouse liver data across multiple datasets and experimental platforms with peak scores occurring during the active phase.

## III. Discussion

In has been found that multiple signaling pathway in mammals are activated rhythmically, according to the daily changes in behavior, nutrient availability, as well as self-sustained circadian clock-driven processes in the individual cells and tissues (Aviram et al., 2021; Goldsmith et al., 2018; Horiguchi et al., 2013; Tsuchiya et al., 2013). However, phosphoproteomics time series data, best suited for pathway activity analyses, is quite limited and thus the transcriptomics datasets remain a major resource for pathway activity quantification. In this study, we have employed gene expression signatures to search for rhythmic “footprints” of pathways activation in published mouse liver transcriptomics datasets.

As the liver transcriptome of wild-type animals is highly rhythmic, with roughly 20% of the genes oscillating with a 24 h period, one might expect that many of the signature genes will be among the rhythmic genes. Indeed, we found that multiple signature genes for various pathways included in PROGENy, are known to be rhythmic in certain tissues, or even in *in vitro* systems, such as Nr1d2 (Rev-Erbβ) in the EGFR signature, or Bhlhe40 (Dec1) in the Hypoxia signature. For mouse liver datasets analyzed here, the final scores for pathway activities also tended to be rhythmic for multiple pathways. Consistent results across datasets were found for 3 pathways: EGFR, PI3K, and p53. It remains to be established, if such rhythmicity in the score correlates well with the rhythmicity in signal transduction through the respective pathways, and what are the delays between the peak activities of the pathways and the peak scores. For the EGFR pathway, we showed which signature genes contribute most to the EGFR score rhythmicity. This analysis revealed a handful of influential genes, whose contributions to the final score were in phase with one another, driving the rhythm in the score.

Previous research of the EGFR pathway provides convincing evidence of the EGFR pathway rhythmicity along the 24 h day, as well as multiple estimates of the peak activity time. Several studies employed immunostaining techniques to evaluate activation of the effector kinases downstream of EGFR, called Erk1 and 2, and produced somewhat conflicting evidence with regard to the peak phases: around CT6 (Chao et al., 2017; Tsuchiya et al., 2013), or CT16 (Lauriola et al., 2014). The lowest levels appeared more consistently at around CT12-14, which has been attributed to the downregulation via glucocorticoids (Lauriola et al., 2014) or circadian clock-controlled phosphatase Mkp1 (Chao et al., 2017). Erk activation is known to lead to its nuclear translocation. However, in the nuclear proteomics study from Wang et al. the nuclear Erk did not show clear rhythms, albeit having the lowest values at CT12 (Wang et al., 2017). A few phosphoproteomics studies quantified Erk activity via the abundance of its phosphorylated targets, and all have estimated the peak activity to happen in the subjective day (around CT6) (Robles et al., 2017; Wang et al., 2017; Wang et al., 2018). These estimates point to a delay of 6-8 hours from a pathway activation to the PROGENy score peak. This can explain the discrepancy in the peak phase estimates acquired from phosphoproteomic data vs. gene expression signature-based scoring.

Regarding PI3K, it has been shown that the peak activation coincides with the peak food intake, which happens for mice in the beginning of subjective night (CT12) (Aviram et al., 2021; Vollmers et al., 2009). Interestingly, a recent dataset from Aviram et al., containing both Western blots and RNA-Seq, allowed us to compare the timing of Akt phosphorylation downstream of PI3K and the PROGENy pipeline-derived peak phase for PI3K (Aviram et al., 2021). For both wildtype mice with a regular 24 h rhythm and mutant mice with a 16 h rhythm in Akt phosphorylation, we also observe a rhythm in PI3K score with a period of 24 and 16 h, respectively. In both cases the delay from the Akt phosphorylation till the score peak is approximately 4 h (Fig.S7).

Previous reports of the rhythmicity in EGFR and PI3K pathways activation, combined with our gene signature analysis, raises an interesting question of the possible sources of such rhythmicity in the signaling network downstream of EGFR and other receptor tyrosine kinases (RTKs). For example, cell autonomous rhythmicity can originate from a crosstalk that has been reported for circadian clock, cell cycle, and the EGFR signaling network (El-Athman et al., 2017; Walker et al., 2007; Wang et al., 2019). Alternatively, there could be a rhythmicity in some inputs to the EGFR network, either local or systemic, that drive the rhythms in the network activation. To gain an overview of various possible sources of daily rhythms in the EGFR pathway, we summarize in Fig.5 some examples of genes and proteins, involved in the signaling downstream of RTKs, that have evidence of rhythmicity, based on Atger transcriptomics data, two liver proteomics datasets, as well as literature (see Table S1 for references). For example, various ligands of EGFR and related receptors have daily rhythms on the transcript level in mouse liver cells. Furthermore, daily rhythms of an EGFR ligand HB-EGF in bloodstream were previously reported. Rhythmicity in the system can also stem from rhythmic levels of adaptor proteins (such as Grb7), from rhythmicity in the regulators of Ras activity, etc. Moreover, other pathways, which exhibit daily rhythms, are involved in crosstalk with RTK signaling. For example, p38 activity has been reported to be rhythmic in the cell culture and is known to regulate EGFR endocytosis.

**Fig.5.**
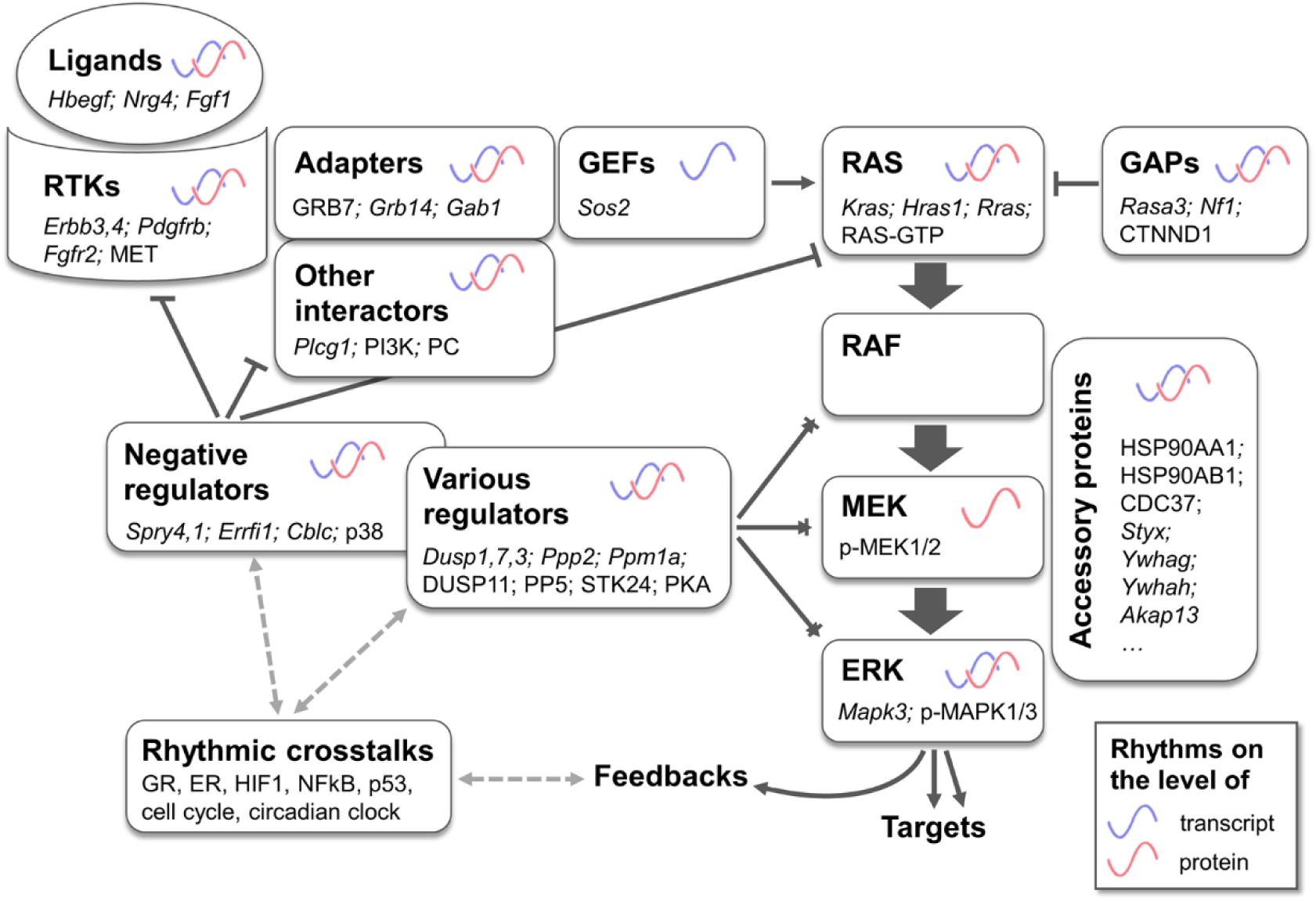
Evidence of rhythmicity in signaling network downstream of EGFR and other receptor tyrosine kinases (RTKs). Reported cell autonomous or non-cell autonomous rhythmicity in the components of EGFR signaling network or their interactors. Rhythmic transcripts are indicated in italics, rhythmic proteins in block letters. Two proteomics (Robles et al., 2014; Wang et al., 2017) and one transcriptomics (Atger et al., 2015) datasets were analyzed for rhythmicity; data for rhythmic posttranslational modifications, membrane lipids, and rhythmic crosstalks stems from literature search (see Suppl. Table 1).

In this project we employed the gene expression signatures to score the activity of various cancer-related pathways in transcriptomics time series data, with a subsequent rhythmicity analysis of the resulting scores. We have found that the highly rhythmic mouse liver data tends to have multiple rhythmic scores of pathway activities, with several pathways displaying consistent patterns across multiple datasets. As healthy cells *in vivo* integrate multiple signaling pathways throughout the day, it remains a challenge to disentangle the individual pathways activation. However, as tumor cells are more reliant on a single dominant pathway for growth and proliferation, a similar gene signature approach applied to tumor samples might assist in evaluating the daily dynamics in the activity of such dominant oncogenic pathways. This in turn can guide chronotherapeutic approaches for different tumor types, i.e. maximizing effectiveness of therapy by means of timed drug administration.

## Methods

### Data processing for PROGENy application

We have selected multiple publicly available mouse liver transcriptomics datasets. In each, mice were entrained by the light-dark cycles and then released into darkness before the beginning of the data collection. RNA-Seq datasets were used in the normalized form, as provided by the authors of respective studies, but without log-transformation to avoid negative values in the data that would cause negative score contribution in PROGENy for positively weighted genes. The microarray data from Zhang et al., acquired with Affymetrix MoGene 1.0 ST arrays, was analyzed with R package *oligo* using RMA normalization. Based on the distribution of intensities, the threshold was set to 4, and only the probes with intensity higher than the threshold in at least 8 out of 24 samples were considered. Probes were cross-referenced to gene symbols with *biomaRt* package. The microarray data from Hughes et al., acquired with Affymetrix Mouse Genome 430 2.0 arrays, was analyzed with R package *affy* using MAS5 normalization and built-in absent/present calling. Only the probes marked as present in 16 out of 48 samples were considered. Values for multiple probes corresponding to one gene were averaged. Both microarray and RNA-Seq data was further normalized via division by the mean expression values, so that every gene has a mean expression value of 1 and amplitudes are relative to the mean.

### Analysis of daily rhythmicity in PROGENy score

The mouse version of PROGENy was applied to the processed time series data to yield time series of pathways activity scores for 14 pathways. The PROGENy parameter *scale* was set to true (scores are normalized to have the average of 0 and the standard deviation of 1). For each pathway, 100 most significant PROGENy signature genes were included in score calculations, the majority of which were present in the dataset, ranging in Atger et al. dataset from 50 signature genes for Trail to 91 for Hypoxia pathway. The resulting score time series (with replicates, when available) were analyzed for presence of rhythmicity with 24 h period with RAIN (default parameter values). The resulting p-values produced by RAIN were adjusted for multiple testing (14 tests) with the Benjamini-Hochberg algorithm, and a threshold of 0.05 was used for resulting q-values to determine the significance of the rhythmicity.

### Jackknife analysis of the individual gene contributions to PROGENy scores

For the “jackknife analysis”, individual signature genes were omitted one at a time, and the score was recalculated. The *scale* parameter was set to false, and instead the scaling was performed in such a way so that a gene removal does not influence the scaling factors: subtracting the mean score of original data (when no gene are removed) and dividing by the standard deviation of the original data.

### Identifying the set of rhythmic transcripts and proteins

RNA-Seq and microarray data were processed as discussed above. The set of rhythmic transcripts was determined with RAIN (FDR < 0.05) and was further filtered to only include the transcripts with mean-to-peak fold-change bigger than 1.2, using the fitted harmonic regression models for the fold-change calculation. For the proteomics data analysis, normalized and log-transformed data was used as is provided by the authors of the respective studies (Robles et al., 2014; Wang et al., 2017). The missing values in the replicates were imputed as averages between all available values among the replicates (for a particular protein at a particular time point). If all replicates at a certain timepoint had missing values, those were not imputed. The set of rhythmic proteins was determined with RAIN (FDR < 0.05) and filtered to have mean-to-peak fold-change bigger than 1.1.

### Selecting set of “influential genes” for EGFR score rhythm

An influential gene was defined as a gene for which relative amplitude multiplied with absolute weight yields a value bigger than a threshold of 1.5 for RNA-Seq datasets, 1.0 for the Hughes et al. microarray dataset (Hughes et al., 2009), or 0.2 for the Zhang et al. dataset (Zhang et al., 2014).

## Acknowledgements

This study was funded by Deutsche Forschungsgemeinschaft (DFG, German Research Foundation), grant RTG2424445 to EW (doctoral training programme CompCancer), grant AN 1553/2-1 to BA, and Project-ID 278001972 – TRR 186 to HH.

## Author contributions

Study idea and design: EW and HH. Bioinformatic analysis: EW, HH and BA. Writing the manuscript (initial version): EW. Editing, reviewing the manuscript: EW, HH, and BA.

