## Supplementary document for "Gene expression signatures predict circadian rhythms in oncogenic pathways"

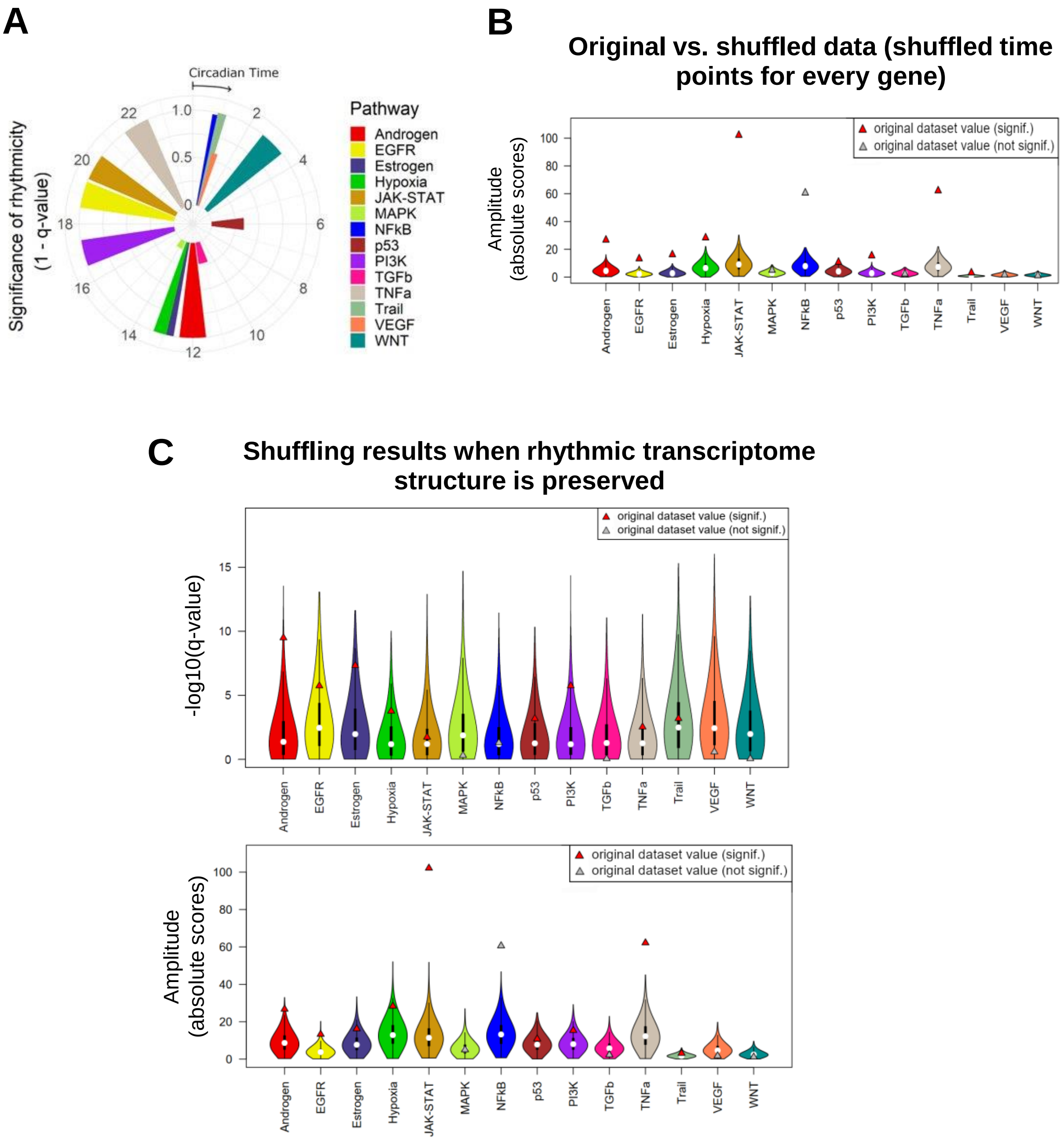


**Fig.S1.** Effects of the normalization step (A,B) and shuffled gene names (C) on the pipeline outcome.

**(A)** The analysis pipeline without normalization step produces results similar to the mail pipeline. Peak phases of PROGENy scores for all 14 pathways are shown; height of the bars corresponds to significance of rhythmicity (1 – q-value) and the dashed red line designates the significance cutoff. **(B)** Distributions of the amplitudes of the score oscillations acquired from shuffled data (times shuffled for each gene) in comparison with original dataset. **(C)** Distributions of the q-values and amplitudes acquired from shuffled Atger et al. dataset (genes names shuffled) in comparison with original q-values. When only gene names are shuffled, the rhythmic structure of the transcriptomics data is preserved and on average 10 PROGENy pathways are rhythmic (q-value above 0.05 cutoff).

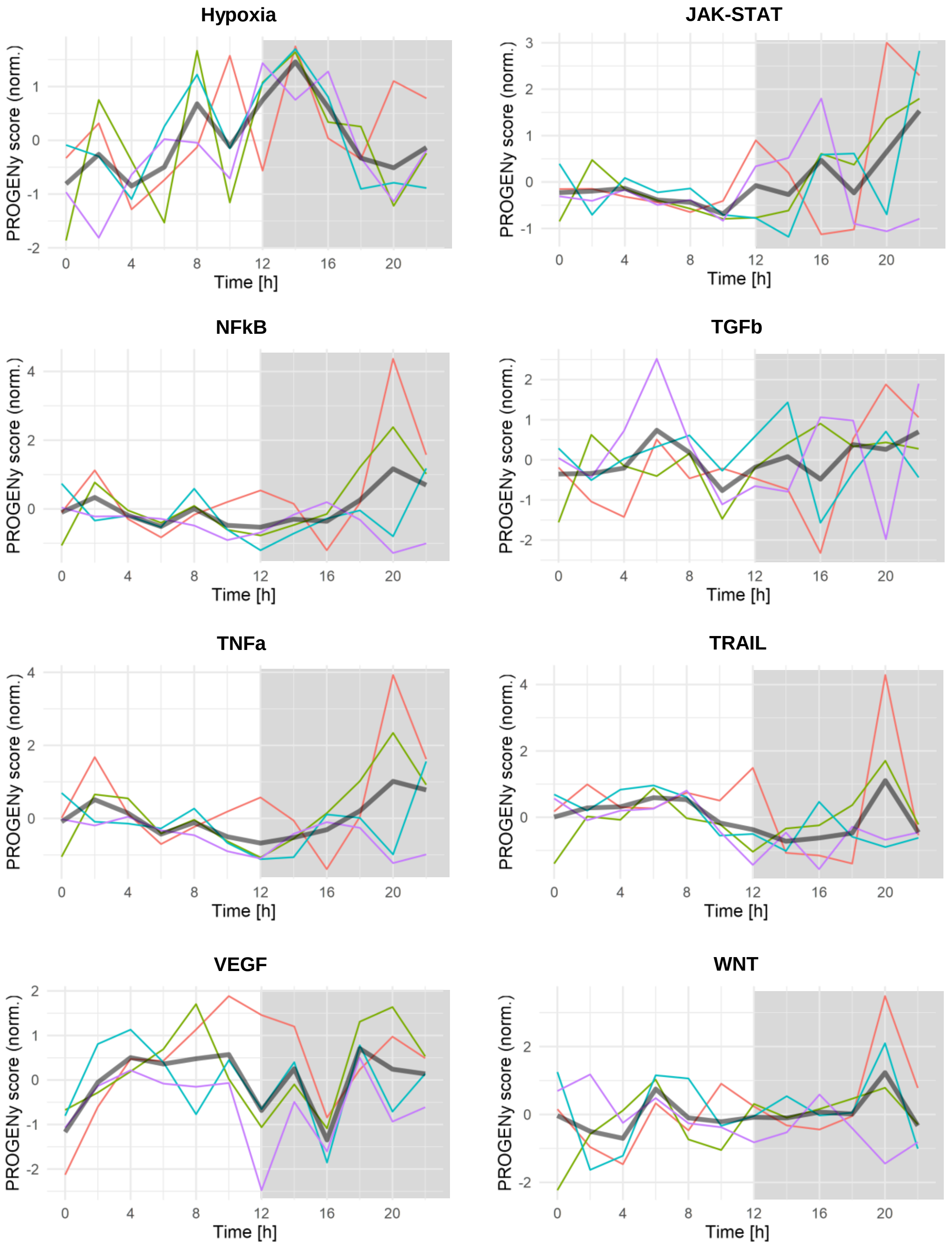


**Fig.S2.** Time series for PROGENy scores for the 8 pathways not shown in Fig.2C.

Colored lines correspond to the individual replicates, labeled as in the Fig2C; grey line shows average. The strong resemblance between NFkB and TNFa is due to a big overlap in their signature gene sets, particularly among most influential genes. Grey lines show averages.

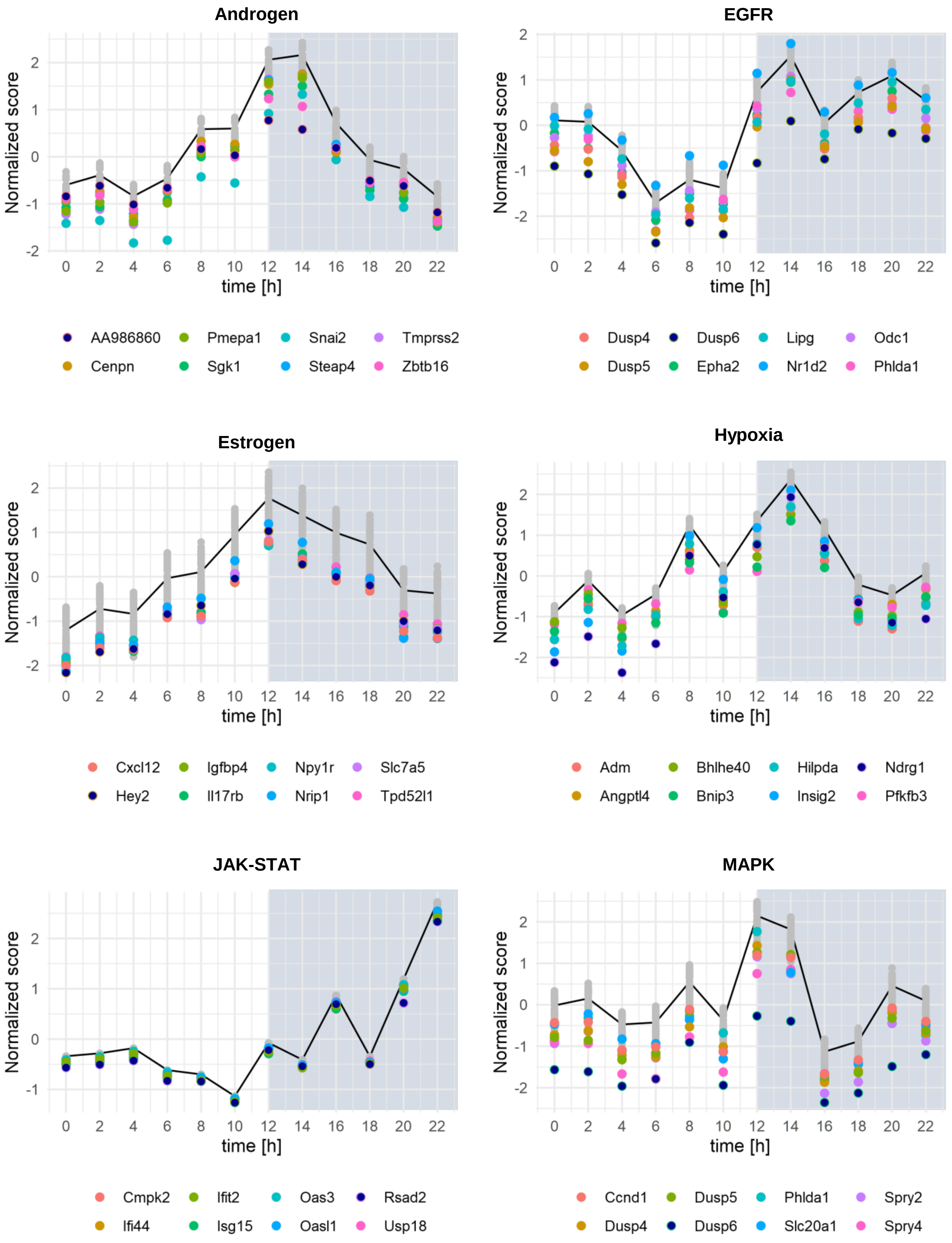

Androgen
Normalized score
time [h]
AA986860
Cenpn
Pmepa1
Sgk1
Snai2
Steap4
Tmprss2
Zbtb16
EGFR
Normalized score
time [h]
Dusp4
Dusp5
Dusp6
Epha2
Lipg
Nr1d2
Odc1
Phlda1
Estrogen
Normalized score
time [h]
Cxcl12
Hey2
Igfbp4
Il17rb
Npy1r
Nrip1
Slc7a5
Tpd52l1
Hypoxia
Normalized score
time [h]
Adm
Angptl4
Bhlhe40
Bnip3
Hilpda
Insig2
Ndrg1
Pfkfb3
JAK-STAT
Normalized score
time [h]
Cmpk2
Ifi44
Ifit2
Isg15
Oas3
Oasl1
Rsad2
Usp18
MAPK
Normalized score
time [h]
Ccnd1
Dusp4
Dusp5
Dusp6
Phlda1
Slc20a1
Spry2
Spry4

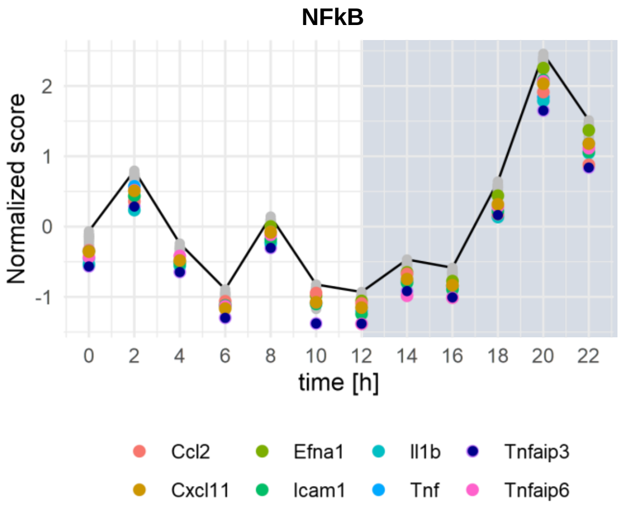


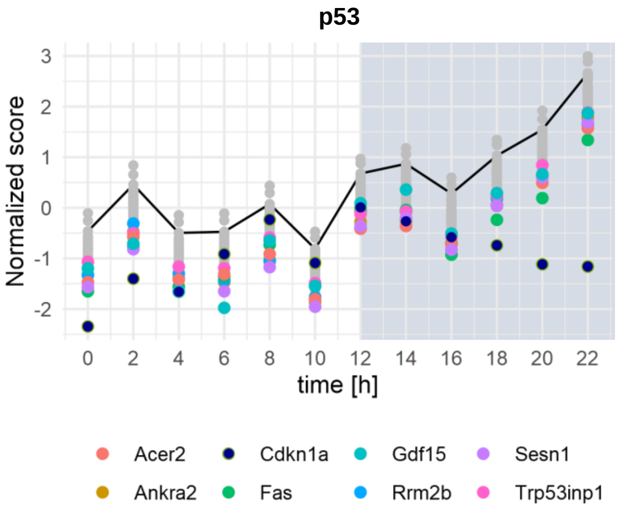


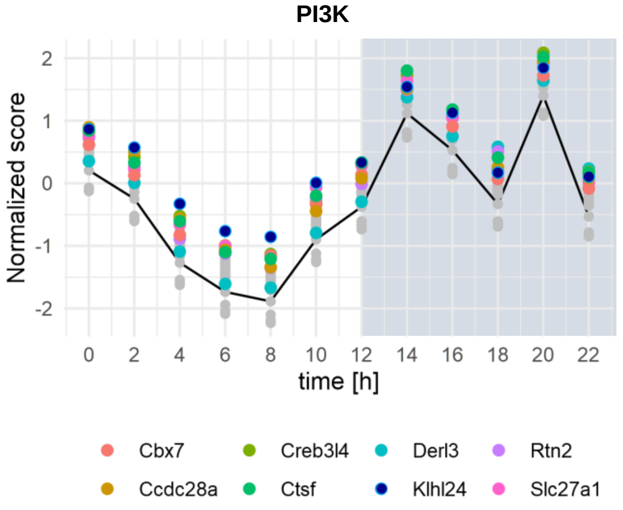


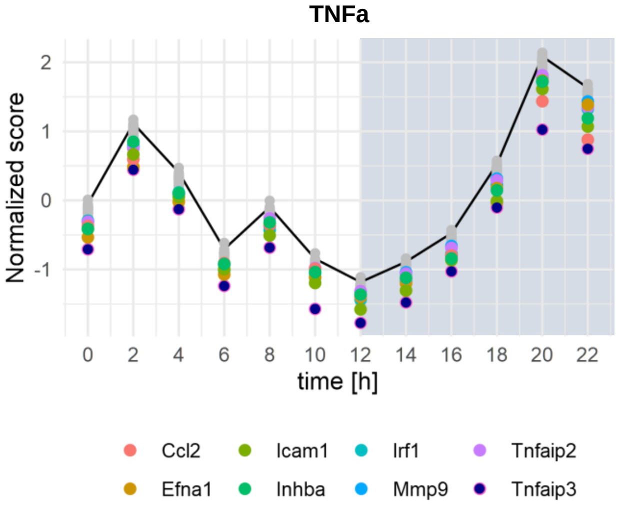


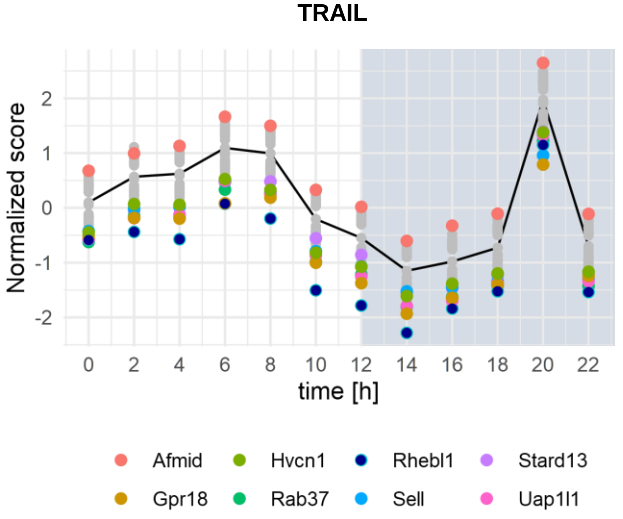


**Fig.S3.** "Jackknife" approach: removal of any one signature gene from the dataset does not abolish rhythmicity of the pathways' activity scores.

Black lines correspond to the time series for PROGENy scores (replicates averaged), when the full dataset is analyzed. Dots represent new scores after removal of one signature gene. Dots for the most influential genes are colored and the names of those genes are indicated. The normalization of the scores (y-axis) is performed so that the original dataset (when no genes are removed) has the mean score of 0, with standard deviation of 1.

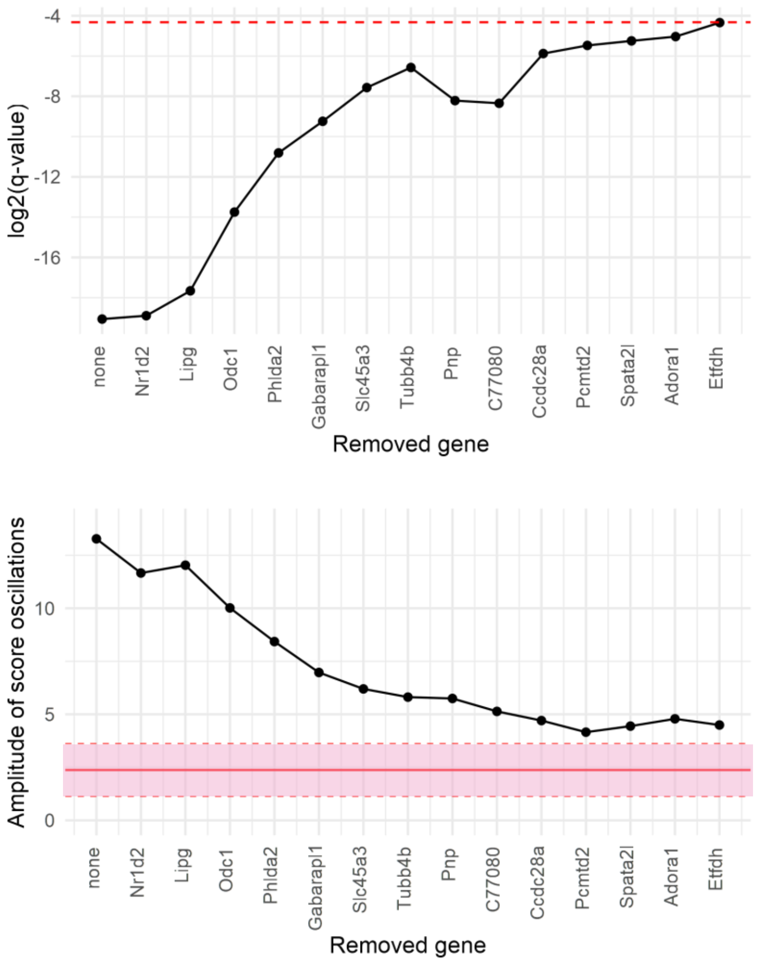


### **Fig.S4.** Contribution of rhythmic signature genes to the rhythmicity of the EGFR score in the Atger dataset.

Rhythmic signature genes were removed from the data in the order of their score contributions (based on their weights and amplitudes) and the EGFR scores were recalculated (replicates averaged). RAIN q-values (top panel) were used to quantify how rhythmic the resulting scores are, and harmonic regression was used to fit an amplitude to the absolute scores (bottom panel). In the top panel, the red dashed line indicates the significance cutoff; in the bottom panel, the solid red line indicates the mean amplitude in the shuffled datasets (where rhythmic structure of the transcriptome was destroyed) and pink shaded areas depict the standard deviation.

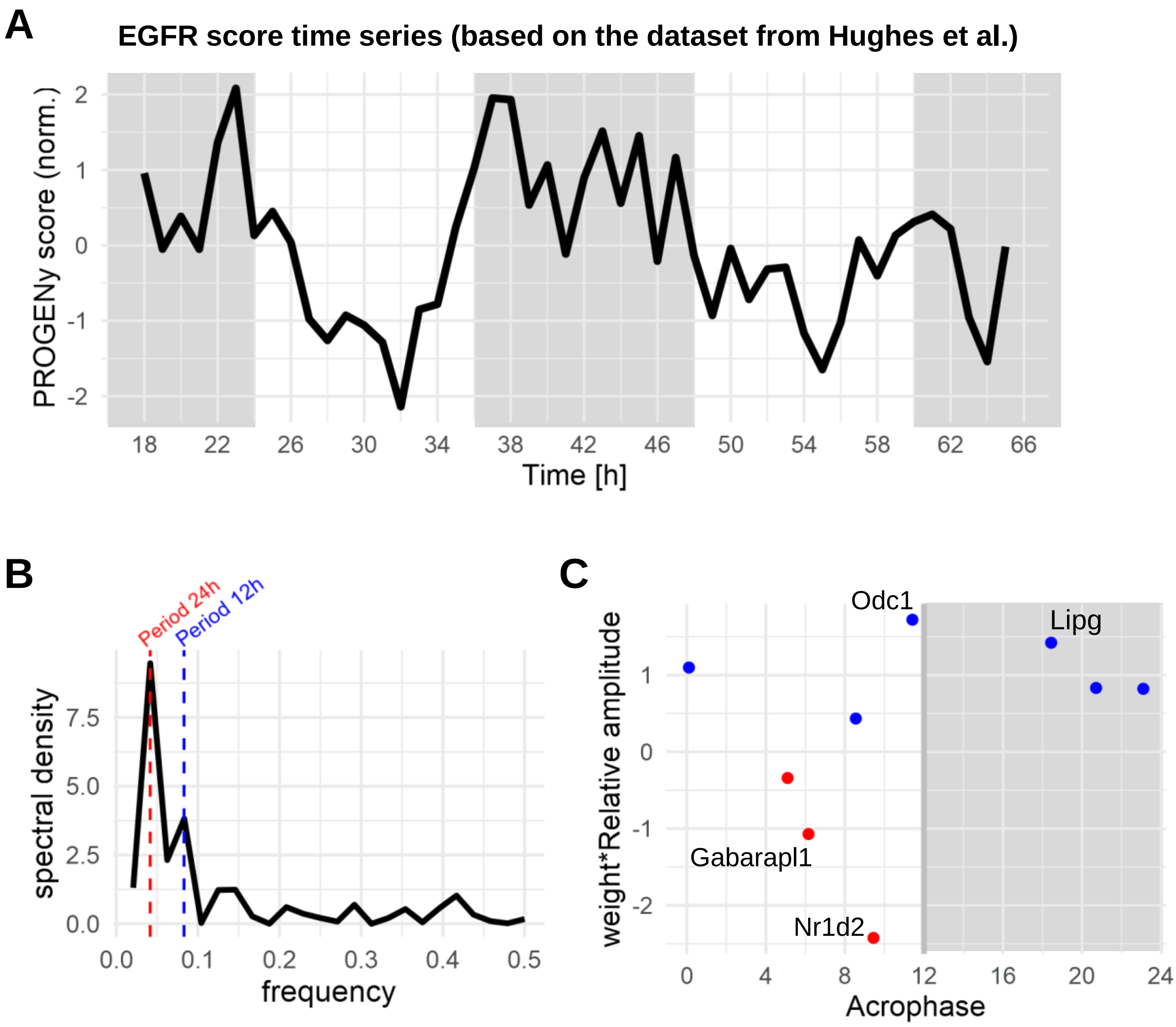


**Fig.S5.** EGFR score has a significant daily rhythm in the Hughes et al. dataset.

**(A)** PROGENy score time series for the EGFR pathway; grey areas on the graph correspond to subjective night hours. **(B)** Spectral analysis of the EGFR score time series using fast Fourier transform and no smoothing reveals the major peak corresponding to a 24 h period, and the secondary peak corresponding to 12 h period. **(C)** Peak phases of rhythmic signature genes plotted against the strength of their contribution to the score, calculated as relative amplitude of gene expression times PROGENy weight; color of the dots corresponds to the sign of the weight coefficient. Genes with most influence on the score are the highest (positive weight) and lowest (negative weight) dots on the graph, their names are indicated.

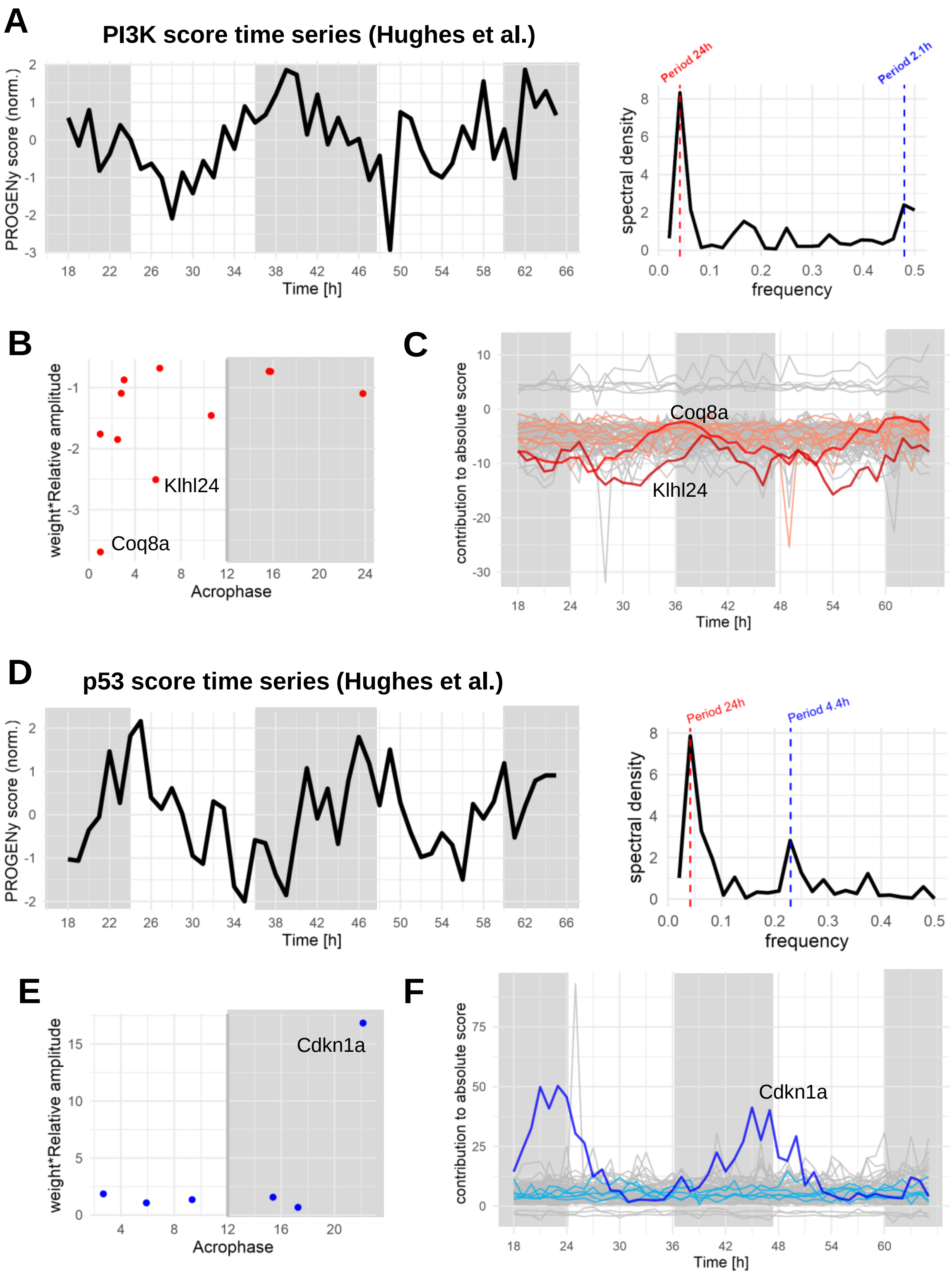


**Fig.S6.** PI3K and p53 scores rhythmicity in the Hughes et al. dataset.

**(A,D)** PROGENy score time series for the PI3K and p53 pathways (left) and corresponding frequency spectra (right); grey areas on the graph correspond to subjective night hours.
**(B,E)** Peak phases of rhythmic signature genes plotted against the strength of their contribution to the score, calculated as relative amplitude of gene expression times PROGENy weight; color of the dots corresponds to the sign of the weight coefficient. Genes with most influence on the score are the highest (positive weight) and lowest (negative weight) dots on the graphs, their names are indicated.
**(C,F)** Contributions to the score from all the signature genes. Rhythmic genes are depicted with colors (blue for positive weight genes, red for negative), and the most influential genes are highlighted with stronger colors; grey areas on the graphs correspond to subjective night hours.

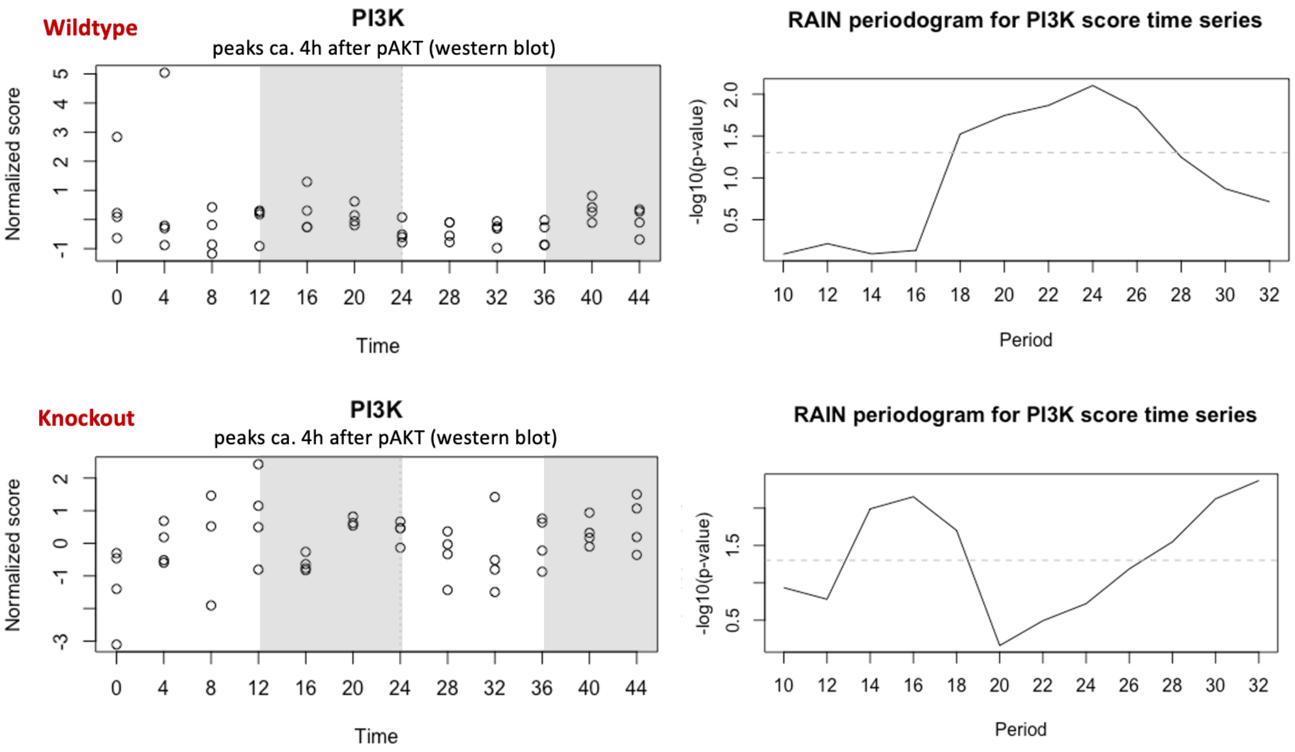


**Fig.S7.** Rhythmicity of the PI3K scores in the Aviram et al. dataset.

Using western blot technique for pAkt, Aviram et al. reported rhythmical activation of PI3K pathway in mouse liver: with 24 h period in wildtype (top panel) and 16 h in Per1/2 double knockout mice (bottom panel). Using RNA-Seq data from this study, we compare these results with the rhythmicity in PI3K PROGENy scores and observe the same periodicity and the delay of approximately 4 h from the pAkt peaks to the PI3K score peaks. Dots represent values from 4 replicates (left subfigures). Periodograms are built analogously to the source study, albeit with RAIN software. Grey areas on the graphs correspond to subjective night hours.

**Table S1.** Rhythmic components of the EGFR network, or rhythmic interactors with the network.

| Name | Function | Type of molecule | Peak time | Dataset / Study reference |
|---|---|---|---|---|
| Hbegf | Growth factor (ligand) | mRNA | CT1, CT4 | [Atger et al., 2015], [Lauriola et al., 2014] |
| HBEGF | Growth factor (ligand) | protein | CT4-10 | Higher levels during the day in blood [Lauriola et al., 2014] |
| Nrg4 (neuregulin) | Growth factor (ligand) | mRNA | CT12 | [Atger et al., 2015] |
| Fgf1 | Growth factor (ligand) | mRNA | CT17 | [Atger et al., 2015] |
| Erbb3 | Growth factor receptor (neuregulins) | mRNA | CT5 | [Atger et al., 2015] |
| Erbb4 | Growth factor receptor (HBEGF) | mRNA | CT8 | [Atger et al., 2015] |
| Pdgfrb | Growth factor receptor | mRNA | CT4 | [Atger et al., 2015] |
| Fgfr2 | Growth factor receptor | mRNA | CT7 | [Atger et al., 2015] |
| MET (HGFR) | Growth factor receptor | protein | CT20 | [Robles et al., 2014] |
| GRB7 | Adaptor | protein | CT4 | [Wang et al., 2017] |
| Grb14 | Adaptor | mRNA | CT4 | [Atger et al., 2015] |
| Gab1 | Adaptor | mRNA | CT11 | [Atger et al., 2015] |
| Plcg1 | Phospholipase C | mRNA | CT10 | [Atger et al., 2015] |
| PC (phospha-tidylcholine) | Cell membrane component | phospholipid | CT22-0 | [Gréchez-Cassiau et al., 2015]; Crosstalk: [Pawar & Sengupta, 2018; Kim et al., 2021] |
| p-AKT | Cell signaling, downstream of PI3K | protein-phosphate | CT12 | [Vollmers et al., 2009; Aviram et al., 2021]; Crosstalk: [reviewed in Mendoza et al., 2011] |
| Sos2 | Ras-GEF | mRNA | CT9 | [Atger et al., 2015] |
| Nf1 | Ras-GAP | mRNA | CT5 | [Atger et al., 2015] |
| Rasa3 | Ras-GAP | mRNA | CT8 | [Atger et al., 2015] |
| CTNND1 (p120) | Ras-GAP | protein | CT23 | [Wang et al., 2017] |
| Kras, Hras1, Rras | GTP-ase, signaling | mRNA | CT21, CT16 | [Atger et al., 2015] |
| RAS-GTP | Active GTP-ase, signaling | protein-GTP | CT6 | [Tsuchiya et al., 2013] |
| p-MEK | Active kinase, signaling | protein-phosphate | CT6 | [Tsuchiya et al., 2013] |
| p-MAPK1/3 (ERK1/2) | Active kinase, signaling | protein-phosphate | CT6 | [Tsuchiya et al., 2013] |
| Mapk3 (Erk1) | Proteinkinase, signaling | mRNA | CT9 | [Atger et al., 2015] |

| Name | Function | Type of molecule | Peak time | Reference |
|---|---|---|---|---|
| Spry4 | Negative regulator of EGFR signaling | mRNA | CT7 | [Atger et al., 2015] |
| Spry1 | Negative regulator of EGFR signaling | mRNA | CT22 | [Atger et al., 2015] |
| Errfi1 | Negative regulator of EGFR signaling | mRNA | CT15 | [Atger et al., 2015] |
| Cblc | E3 ubiquitin ligase, EGFR degradation | mRNA | CT7 | [Atger et al., 2015] |
| p-p38 | Cell signaling; regulates endocytosis of EGFR | protein-phosphate | CT12-16 | [Goldsmith et al., 2018]; Crosstalk: [Frey et al., 2006; Verdaguer et al., 2021] |
| Dusp1,7,3 | Protein phosphatases, negative regulators | mRNA | CT14-16 | [Atger et al., 2015] |
| DUSP11 | Protein phosphatase | protein | CT18 | [Wang et al., 2017] |
| Ppp2r2d, Ppm1a | Ser/Thr phosphatases, negative regulators | mRNA | CT7, CT10 | [Atger et al., 2015] |
| PPP5C | Ser/Thr phosphatase, negative regulator | protein | CT23 | [Wang et al., 2017] |
| STK24 | Ser/Thr protein kinase, upstream of MAPKs | protein | CT2 | [Wang et al., 2017] |
| Prkar2a; and other subunits | Protein kinase A, can act as a positive or negative regulator | mRNA | CT20; CT0-3 | [Atger et al., 2015];<br>[Yan et al., 2008] |
| HSP90AA1 ,AB1 | Chaperone, accessory protein | protein | CT23 | [Wang et al., 2017] |
| CDC37 | Co-chaperone to Hsp90 | protein | CT23 | [Wang et al., 2017] |
| Styx | Pseudophosphatase | mRNA | CT10 | [Atger et al., 2015] |
| Ywhag, Ywhah | 14-3-3 family chaperone | mRNA | CT17 | [Atger et al., 2015] |
| Akap13 | scaffold protein, links cAMP and EGFR signaling | mRNA | CT12 | [Atger et al., 2015] |
| nuclear GR | Glucocorticoid receptor | protein | CT12-14 | [Wang et al., 2017];<br>Crosstalk: [Lauriola et al., 2014] |
| Esr1; nuclear ER | Estrogen receptor α | mRNA; protein | CT18; CT0; | [Atger et al., 2015; Wang et al., 2017];<br>Crosstalk: [Britton et al., 2006] |
| Nuclear HIF1 | Active HIF1 | protein | CT8-12 | [Adamovich et al., 2017];<br>Crosstalk: [Miyamoto et al., 2015] |
| Nfkbia | Component of NF-κB pathway | mRNA | CT9 | [Atger et al., 2015];<br>Rhythm in the pathway activation: [Spengler et al., 2012];<br>Crosstalk: [Shostak & Chariot, 2015] |
| Cdkn1a (p21) | Component of p53 pathway | mRNA | CT21 | [Atger et al., 2015];<br>Crosstalk: [Volman et al., 2022] |
